# Repurposing AZD-5991 for inhibiting growth and biofilm formation of *Staphylococcus aureus* by disrupting the cell membrane and targeting FabI

**DOI:** 10.1101/2025.02.13.638150

**Authors:** Yuanyuan Tang, Han Deng, Zhichao Xu, Zhijian Yu, Yong Xiang, Zewen Wen, Shiqing Han, Zhong Chen, Tieying Hou

## Abstract

*Staphylococcus aureus* infections have already become a major threat to public health worldwide. Drug resistance and biofilm formation are two critical factors that significantly affect the efficacy of the antimicrobial treatment of *S. aureus* infections using conventional antibiotics. Therefore, the discovery of novel antimicrobial agents with potent antibacterial and antibiofilm activity has become a hotspot in recent years. Here, we first demonstrated the remarkable inhibitory activity of AZD-5991, a selective Mcl-1 inhibitor, against *S. aureus*. The MIC_50_ and MIC_90_ of AZD-5991 against *S. aureus* were 12.5 μM, and AZD-5991 significantly inhibited the growth of *S. aureus* at a subinhibitory concentration of 1/2 × MIC. Besides, AZD-5991 displayed bactericidal activity and a robust capacity for inhibiting biofilm formation against *S. aureus* with very low drug toxicity against host cell lines. Our data demonstrated the disruption of *S.aureus* cell integrity by AZD-5991 through membrane permeability assays and the ingredients of the bacterial phospholipids could neutralize the antibacterial activity of AZD-5991. Moreover, whole-genome sequencing and proteomic analysis were also applied to gain insights into the possible impact of AZD-5991 on the fatty metabolism of *S. aureus*. Furthermore, the antibacterial activity of AZD-5991 was significantly elevated by exogenous fatty acids linoleic acid (C18:2 Δ 9,12) and arachidonic acid (C20:4 Δ 5,8,11,14). Lastly, the biolayer interferometry assay supported the direct interaction of AZD-5991 with FabI, a necessary protein related closely to bacterial growth and fatty acid metabolism. Conclusively, this study suggests that AZD-5991 could inhibit the planktonic growth and biofilm formation of *S. aureus* by disrupting the cell membrane and targeting FabI. AZD-5991 might be a promising new antibiotic candidate for the antimicrobial treatment of *S. aureus* infections resistant to traditional clinical drugs.

**Graphical Abstract:** 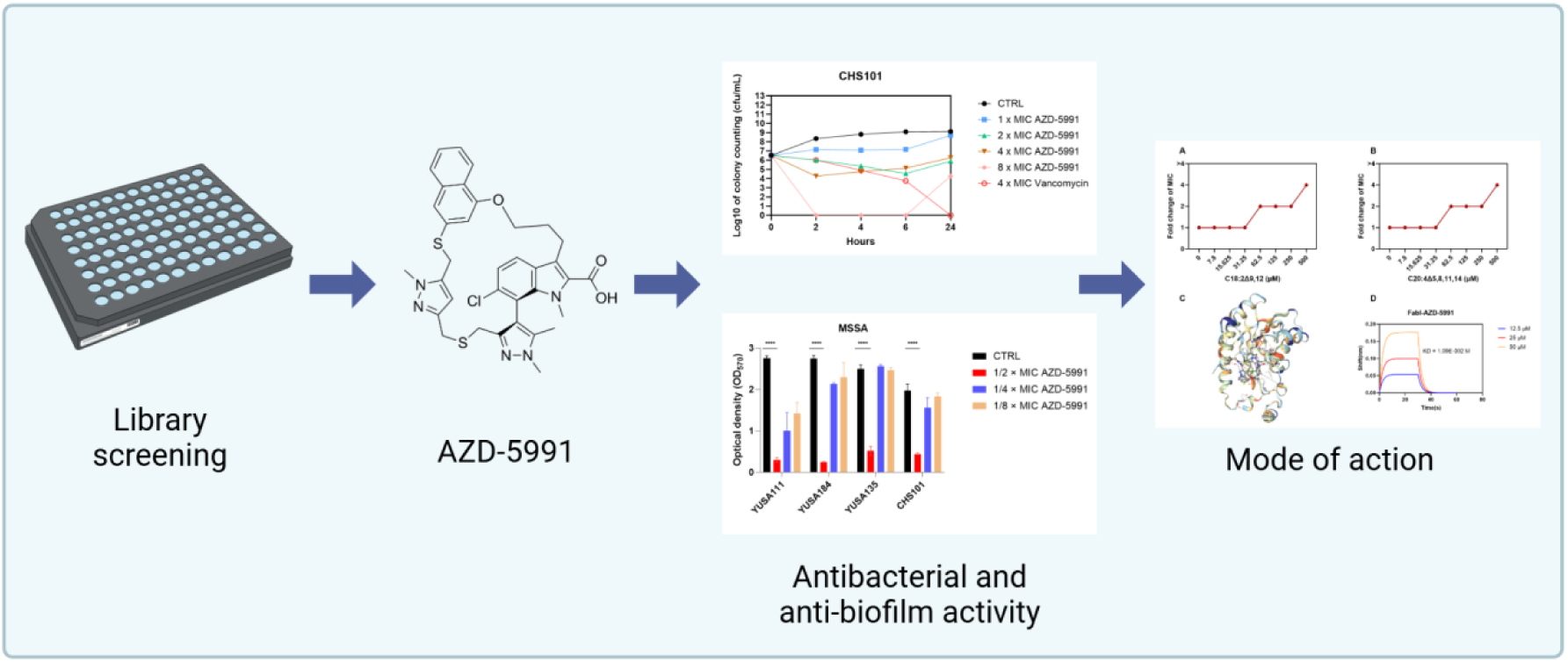

## Introduction

*Staphylococcus aureus* is one of the most well-known species of the Staphylococcus genus, which is frequently seen in a variety of infectious disorders that are acquired in hospitals or the community, such as pneumonia, endocarditis, abdominal infection, skin and soft tissue infection, and more [1]. In recent years, the emergence of *S. aureus* with multi-drug resistance, including methicillin-resistant *S. aureus* (MRSA), linezolid-resistant *S. aureus*, daptomycin-nonsusceptible *S. aureus*, gravely threatening social and public health [2]. According to the surveillance results from CHINET in 2021, the detection frequency of MRSA from clinical samples in China was 30.0% [3]. Linezolid-resistant and daptomycin-nonsusceptible *S. aureus* have been gradually reported in clinics. The high isolation frequency of MRSA and the dissemination of multi-drug *S. aureus* isolates significantly narrow the therapeutic choices for improving the clinical outcome of *S. aureus* infection [4].

Moreover, the formation of biofilms is also one of the major mechanisms by which *S. aureus* evades the host immune eradication and pushes the gradual development of bacterial cells toward the drug-resistance phenotype [5]. The aggregation, proliferation, and adhereance of *S. aureus* cells on some special devices or surfaces can secrete the extracellular polymer molecules and promote biofilm formation. Biofilm often can be found with the formation of viscous and distinct three-dimensional extracellular matrix of *S. aureus* cells. Therefore, *S. aureus* are embedded in biofilm and the biofilm clings to them to create a barrier between the outer environment and bacterial cells [6]. The consequence of biofilm formation is that antibiotics can’t interact directly with biofilm-embed bacterial cells. Till now, just a limited several antibiotics can be used for antimicrobial treatment of biofilm-related *S. aureus* infection [7]. Therefore, one of the current research hotspots is the development of novel antimicrobial agents for treating *S. aureus* infections with drug resistance and biofilm formation [8].

AZD-5991 has been developed as a selective Mcl-1 inhibitor for many years and has been approved by Food and Drug Administration (FDA) for a clinical trial for treating hematological carcinoma in Phase I [9]. Here, we first discovered the antibacterial activity of AZD-5991. The purpose of this study was to examine the antibacterial and anti-biofilm properties of AZD-599 against *S. aureus* in vitro using the growth curve and bactericidal curve. The action mechanism of AZD-5991 on *S. aureus* was preliminarily investigated through membrane proton kinetic potential, the proteome of planktonic *S. aureus*, and the binding experiment of the potential target. This study will provide a basis for new applications of AZD-5991 as the development of antibacterial drugs.

## Result

### AZD-5991 exhibits the robust bacteriocidal activity against Gram-positive bacteria

The MIC of AZD-5991 against Gram-positive bacteria, including *S. aureus*, *S. epidermitis*, *E. faecalis*, and *E. faecium*, was measured by agar dilution, suggesting its range from 6.25 μM to 25 μM (Table 1). The specific MIC values for various bacteria can be found in the supplementary materials (Table S1). In addition, the inhibition activity of AZD-5991 on the planktonic growth of MRSA and *E. faecalis* was further investigated by the bacteria automatic growth curve experiment, further demonstrating AZD-5991 with the concentration of 1 × MIC could completely inhibit the planktonic growth of MRSA YUSA139 and YUSA145 (Figure 1A and 1B), and *E. faecalis* 16C30 and 16C51 (Figure 1C and 1D), respectively. Whereas, AZD-5991 at a concentration of 1/2 × MIC could just inhibit the planktonic growth of YUSA139 and YUSA145 at the early phase of drug exposure and have no impact on the the bacterial growth at 24h.

**Figure 1.**
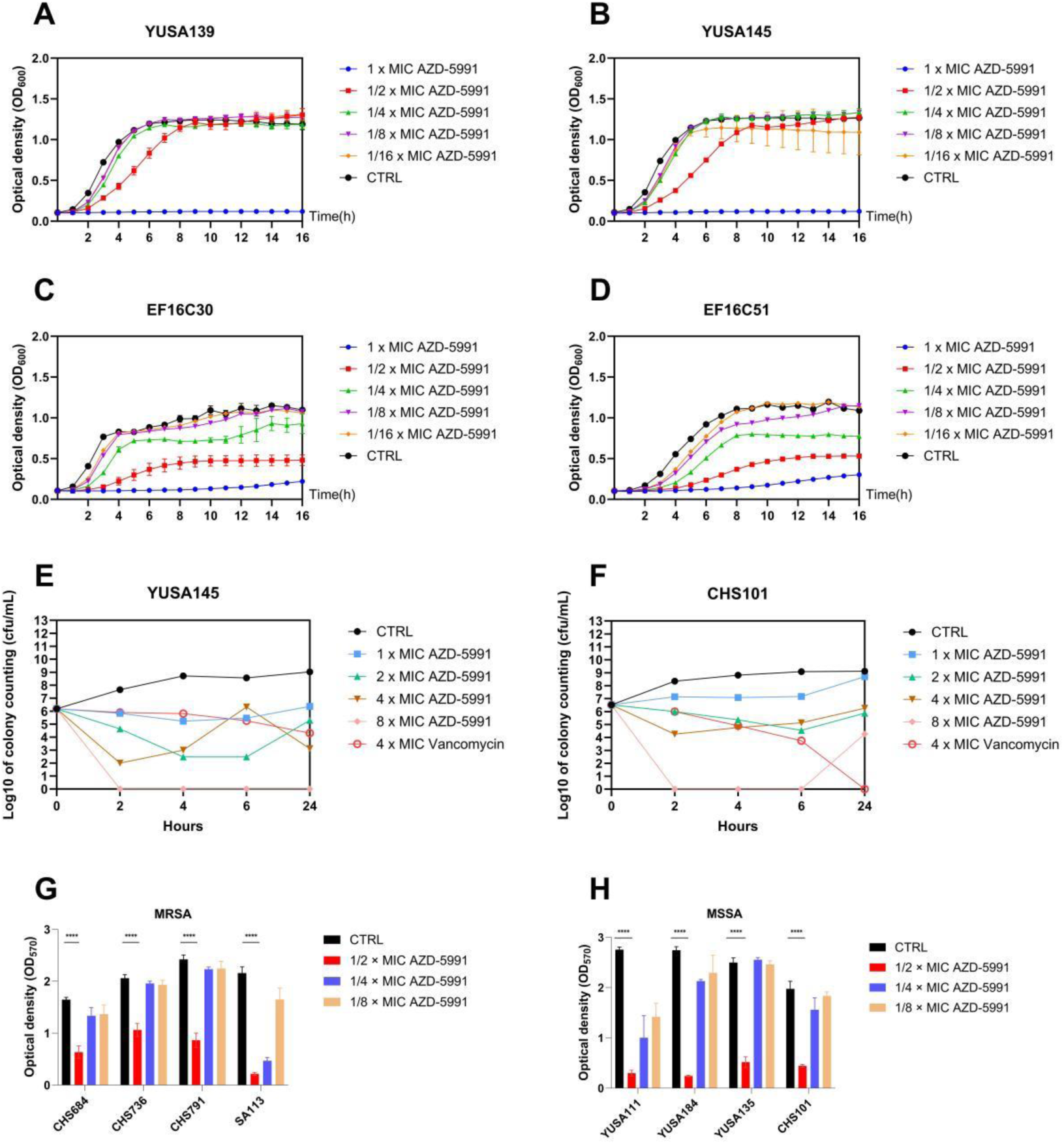
AZD-5991 significantly inhibited the growth of Gram-positive bacteria and the biofilm formation of *S*. *aureus*. (A and B) The planktonic cells of MRSA YUSA139 and YUSA145, and (C and D) *E. faecalis* 16C30 and 16C51 were treated with 1/16 ×, 1/8 ×, 1/4 ×, 1 × MIC of AZD-5991. The OD_600_ value of bacterial cells was measured by Bioscreen C (Turku, Finland) at 1-h intervals for 16 h. TSB without antimicrobials was used as a negative control. Data are shown as Mean ± SEM (n = 3). All experiments were performed in triplicate. (E) *S*. *aureus* YUSA145 and (F) CHS101 were challenged with 1 ×, 2 ×, 4 ×, 8 × MIC of AZD-5991 as well as 4 × MIC of vancomycin. Bacteria growth in TSB without antimicrobials was used as an untreated control. Data are representative of three independent experiments. (G-H) The significant inhibition of AZD-5991 on the biofilm formation of MRSA and MSSA.

**Table 1.** MICs distribution of AZD-5991 against Gram-positive bacterial isolates.

| Bacteria species | AZD-5991 MIC values (μM) | | | MIC<sub>50</sub>/MIC<sub>90</sub> |
| --- | --- | --- | --- | --- |
|  | 6.25 | 12.5 | 25 |  |
| *MRSA* (n=20) | 6 | 14 | - | 12.5/12.5 |
| *MSSA* (n=17) | 2 | 15 | - | 12.5/12.5 |
| *S. epidermids* (n=21) | 6 | 15 | 3 | 12.5/12.5 |
| *E. faecium* (n=19) | - | 10 | 9 | 12.5/25 |
| *E. Faecalis* (n=20) | - | 3 | 17 | 25/25 |

MIC<sub>50</sub>/MIC<sub>90</sub>, the MIC values for 50% or 90% of bacterial growth inhibition.

In order to investigate the antibacterial activity of AZD-5991 in a dose-dependent manner, the time-kill assay of AZD-5991 against *S. aureus* was conducted in comparison to vancomycin. As shown in Figure 1E, The time-kill curve experiment suggested that the bactericidal activity of AZD-5991 with 1 × MIC in YUSA145, was similar to 4 × MIC of vancomycin and in CHS101, that with 4 × MIC was similar to 4 × MIC of vancomycin (Figure 1F). In addition, AZD-5991 with 8 × MIC exhibited a powerful eradication effect on YUSA145 at 4h after drug exposure, suggesting the rapid and efficient bactericidal activity of AZD-5991 against planktonic bacteria.

### The anti-biofilm activity of AZD-5991 against *S. aureus*

The impact of AZD-5991 on the biofilm formation of *S. aureus* was studied using *S. aureus* clinical isolates including MSSA and MRSA by crystal violet staining. The anti-biofilm activity of AZD-5991 with sub-MIC (1/8 ×, 1/4 ×, 1/2 × MIC) could significantly inhibit the biofilm formation of *S. aureus* in a dose-dependent manner, especially at 1/2 × MIC, the biofilm formation in the majority of *S. aureus* clinical isolates could be decreased approximately by 80% (Figure 1G and Figure 1H), indicating that AZD-5991 could significantly inhibit the biofilm formation of *S. aureus*.

### Cytotoxicity of AZD-5991 against the host cells

The cytotoxicity of AZD-5991 was evaluated with CCK-8 assays in different cell lines, including LX-2, HepG2, HCT116, A549 and 293T. No significant cytotoxicity was observed in the LX-2 cell line incubated with AZD-5991 at concentrations ranging from 0 to 50 µM for 24 h, while notable cytotoxicity was detected when the concentrations of AZD-5991 exceeded 100 µM (Figure S1A). The IC_50_ value of AZD-5991 to HepG2 and A549 cells obtained from CCK-8 assay was 50 μM, whereas that were 25 μM for both HCT116 and 293T cells (Figure S1B-E).

### Proteomic analysis of *S. aureus* treated with AZD-5991

Quantitative label-free proteomic analysis investigated the global proteomic response of *S*. *aureus* treated with AZD-5991 during the exponential growth phase. Overall, 1,379 proteins were confidently identified (matched peptides ≥ 1, and FDR <0.01) and quantified, 279 proteins with significantly different expression levels (≥|2|-fold change, p≤0.05), containing 95 up-regulated and 184 down-regulated proteins in AZD-5991 treatment, were determined when compared with the control group (Figure 2A and 2B). Table S2 lists the proteins that are significantly downregulated or upregulated. Detailed information on the functional proteins with significantly different expression levels have been listed in Table S2. In addition, functional analysis of proteins with differentially expressed levels was conducted according to the gene ontology (GO) terms of biological process, molecular function, and cellular component (Figure 2C), suggesting the function proteins with different expression levels in AZD-5991-treated *S. aureus* can be classified to the hydrogen ion transport transporter activity and alpha-amino acid biodynamic process.

**Figure 2.**
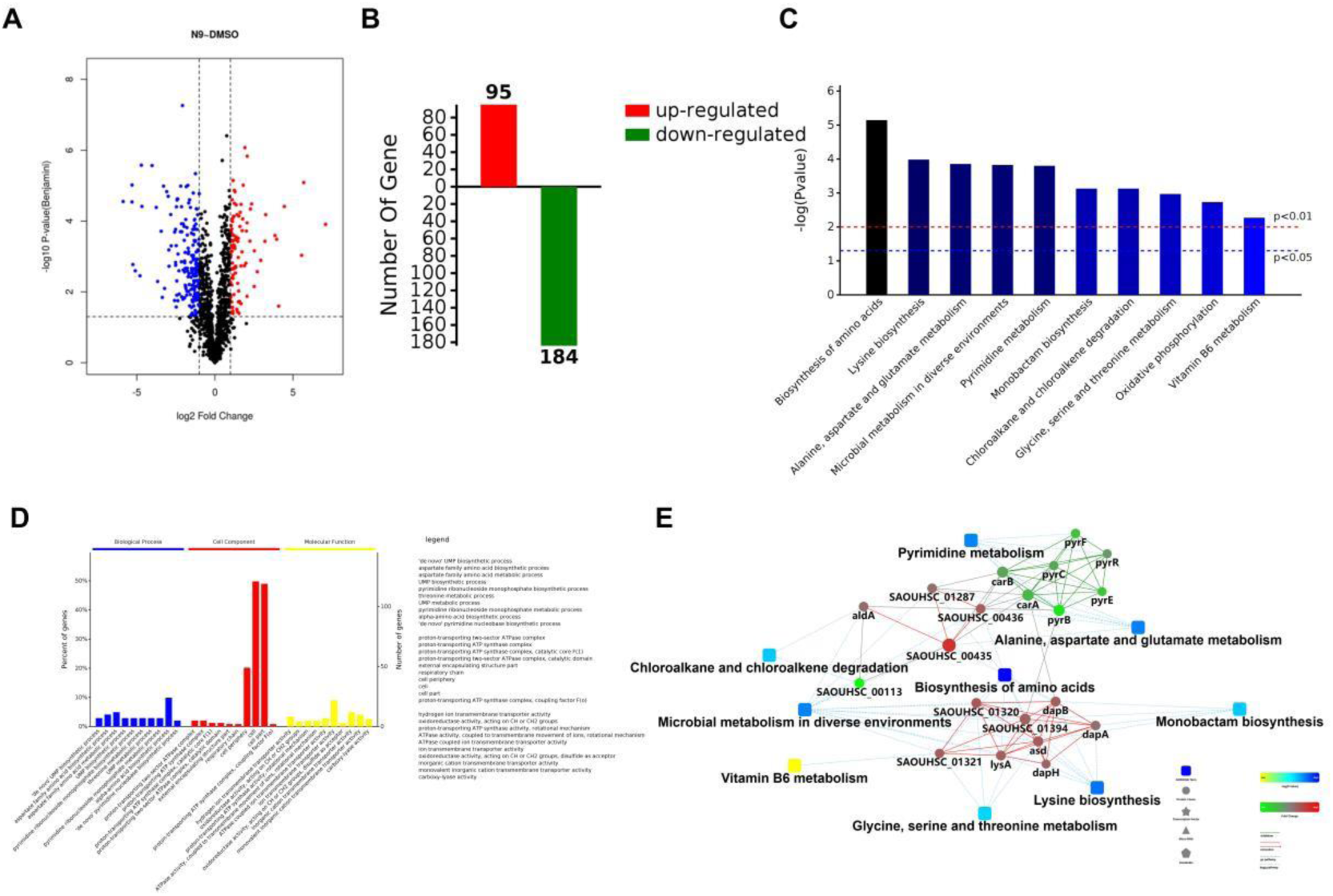
Differentially expressed proteins between the control groups and AZD-5991-treated groups. (A-B) Volcano plots and statistical analysis showed a log2 fold change of protein levels after treatment of *S*. *aureus* SA113 cells with AZD-5991 (1/2 × MIC) compared to DMSO treatment. Blue dots represent proteins whose expression was inhibited by AZD-5991. Red dots represent proteins that are up-regulated by AZD-5991. Data represent average values and p-values were calculated using a two-sided two-sample t-test; n= 3 independent experiments per group. (C) GO annotation and (D) KEGG analysis of the differentially expressed proteins between two groups. (E) Protein-protein interaction (PPI) networks were used to analyze the representative proteins and signal transduction pathways affected by 1/2 × MIC of AZD-5991 treatment in *S*. *aureus* SA113. Proteins down-regulated or up-regulated by AZD-5991 in *S*. *aureus* are marked in green or red, respectively.

Moreover, KEGG analysis showed that the metabolic pathways involved in the proteins with differential expression levels were enriched in the biosynthesis of amino acids, lysine biosynthesis, alanine, aspartate and glutamate metabolism, microbial metabolism in diverse environments, and pyrimidine metabolism pathway (Figure 2D). Protein-protein interaction (PPI) network analysis was constructed using the STRING database (Figure 2E). The results suggested that interactions among downregulated proteins primarily centered around delta-hemolysin hld, L-threonine dehydratase catabolic tdcB, aspartate carbamoyltransferase pyrB, alanine dehydrogenase 1, bifunctional enzyme pyrF/pyrE, immunoglobulin-binding protein spa and sbi as well as ferric uptake regulation protein. These proteins mainly involved in metabolic pathways associated with transferase activity and amino acid transmembrane transport. Upregulated proteins were enriched in ketol-acid reductoisomerase ilvC, large ribosomal subunit protein rpmC, acylphosphatase, ferredoxin katA, and aspartate-semialdehyde dehydrogenase. These proteins primarily participate in metabolic pathways associated with translation and the hydrogen peroxide catabolic process.

### Determination of genetic mutation in AZD-5991-treated *S. aureus* by whole-genome sequencing

Following 40 generations of continuous passage of YUSA145 under AZD-5991 pressure in vitro, the AZD-5991-induced resistant *S. aureus*, called YUSA145N40, was selected and its MIC of AZD-5991 was raised with 8-fold elevation to 100 μM (Figure 3). The genetic mutations in the AZD-5991-induced tolerant *S. aureus* YUSA145N40 were detected by whole-genome sequencing, suggesting three nonsynonymous mutations were found in 3 function genes, including the gene encoding fatty acid efflux pump transcriptional regulator FarR, the gene encoding helix-turn-helix domain-containing protein and the gene encoding threonylcarbamoyl-AMP synthase. Fatty acid efflux pump transcriptional regulator FarR plays a critical role in the metabolism of long-chain fatty acids (Table 2). We thus hypothesized that AZD-5991 exposure might impact cell membrane synthesis and fatty acids metabolism pathway.

**Figure 3.**
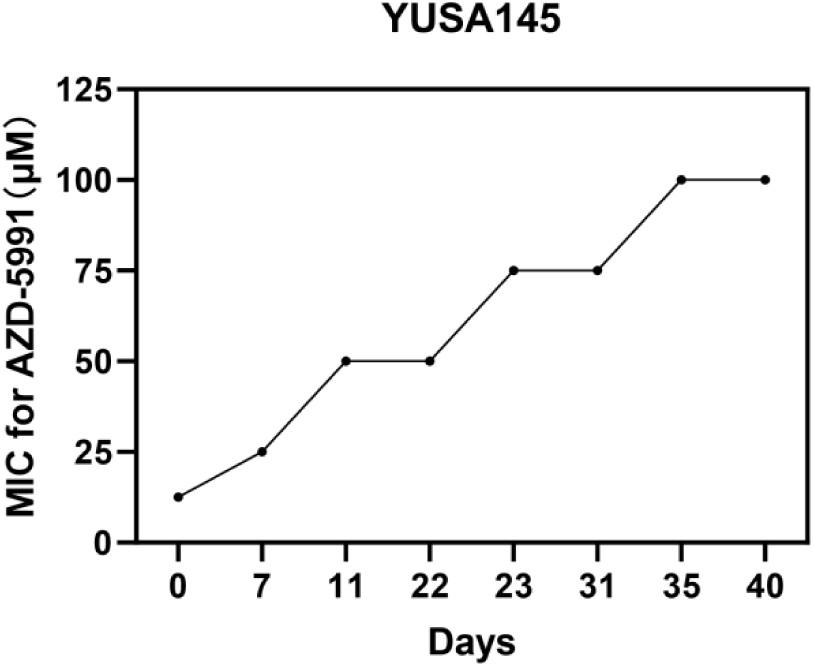
The dynamics of AZD-5991 MIC values in AZD-5991-induced *S*. aureus YUSA145.

**Table 2.**
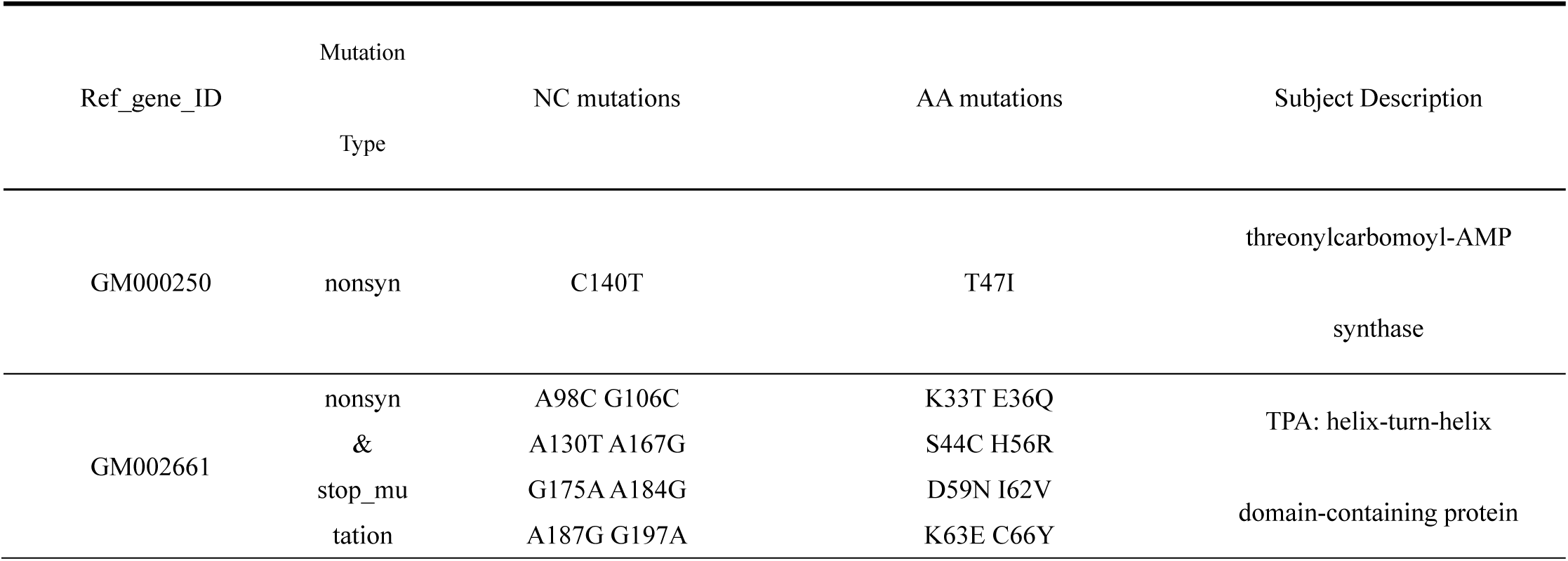
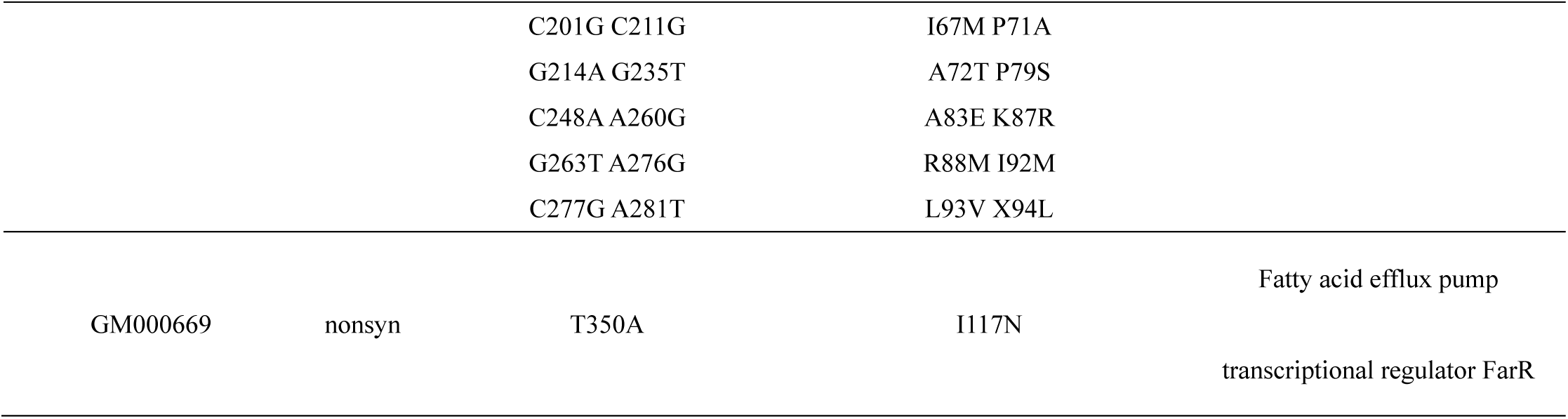
Determination of genetic mutations in YUSA145N40 by whole-genome sequencing.

## Disruption of bacterial cell membranes permeability by AZD-5991

To determine the impact of AZD-5991 on the cell membrane of *S*. *aureus*, the destructive effect of AZD-5991 on cell membrane depolarization and membrane permeability could be assessed by DiSC3(5) and Propidium Iodide (PI) dye assay, respectively. The fluorescence intensity in *S. aureus* SA113 by DiSC3(5) staining can be used for evaluating the depolarizing effect of chemicals on the cell membrane. Our data indicated that under exposure with concentrations of 1 × MIC or 2 × MIC, AZD-5991 exhibited a depolarizing effect on the cell membrane, similar to that of 0.1% Triton X-100 (Figure 4A). Moreover, the fluorescence intensity of PI under AZD-5991 exposure with 4 × MIC in *S. aureus* SA113 was immediately increased to 1.2-fold compared with the control group (Figure 4B). In general, our data solidly indicated the rapid disruption effect of AZD-5991 on the membrane permeability against *S. aureus* cells in a dose-dependent manner.

**Figure 4.**
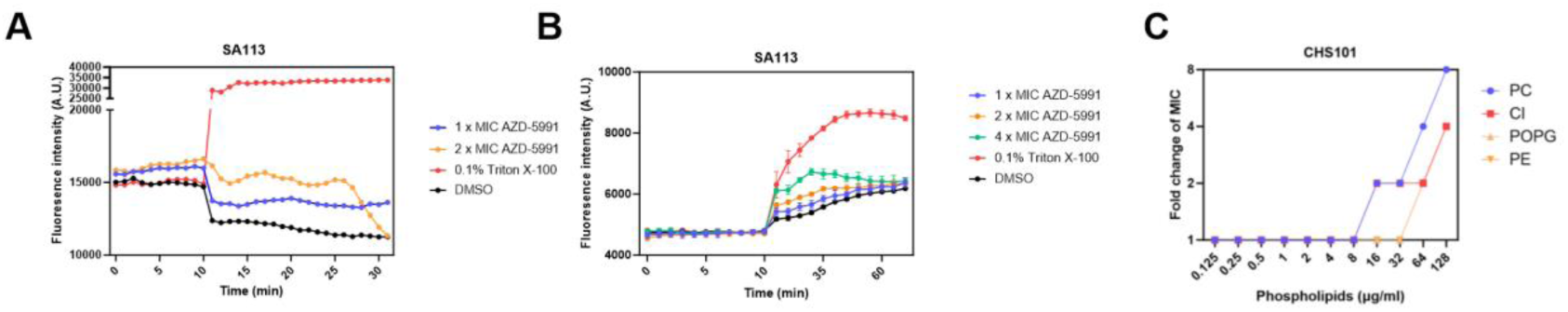
The effect of AZD-5991 on the disruption of bacterial cytoplasmic membrane. (A-B) Membrane depolarization (A) and permeability (B) by AZD-5991 against *S*. *aureus* SA113. (C) Membrane phospholipid neutralization experiment of *S*. *aureus* CHS101.

Membrane phospholipids, including phosphatidylcholine (PC) and cardiolipin (CL), are the major ingredients of *S. aureus* cell membranes. Several reports have demonstrated that the antibacterial activity of multiple antimicrobial agents might be neutralized by PC or CL [10–11]. Therefore, to further explore the antibacterial mechanism of AZD-5991 against S. aureus, four kinds of membrane phospholipids, including PC, CL, POPG, and PE, were exogenously added into bacterial culture medium and further observed whether they could affect the MIC value of AZD-5991 in *S. aureus* CHS101. Our result showed that the MIC value of AZD-5991 in *S. aureus* was significantly increased to more than 2-fold after adding PC and CL with concentrations greater than or equal to 16 μg/ml (Figure 4C). However, no binding effect between AZD-5991 and cardiolipin was found by the BLI kinetic analysis in this study (data not shown).

### AZD-5991 impacts fatty acids pathway by targeting FabI

Previous results demonstrated the potential critical role of the fatty acid pathway in the inhibition of AZD-5991 against *S. aureus*. Here, the impact of exogenous fatty acid components in the culture medium with different concentrations on the antibacterial activity of AZD-5991 was examined and our data from the checkerboard assay (Figure 5A and 5B) demonstrated that the unsaturated long-chain fatty acids linoleic acid and arachidonic acid could significantly increase MIC values and reduce the antibacterial activity of AZD-5991 in a concentration-dependent manner against *S. aureus*. The binding potential of FabI with AZD-5991 was firstly predicted through molecular docking studies. The results indicated that AZD-5991 could bind to FabI in a favorable conformation, yielding a low binding energy of −8.3 kcal/mol. A smaller score value (greater absolute value of the negative value) signifies a stronger binding force, and an affinity that is lower than 7 kcal/mol suggests a good binding force. The best conformation of the AZD-5991-FabI complex is showed in Figure 5C. These findings demonstrate that AZD-5991 has the potential to bind effectively with FabI. Furthermore, the results of the BLI kinetic study supported the direct binding between AZD-5991 and FabI (Figure 5D). Therefore, we propose that AZD-5991 might exert its antibacterial activity by affecting the fatty acid-related pathways in *S*. *aureus*.

**Figure 5.**
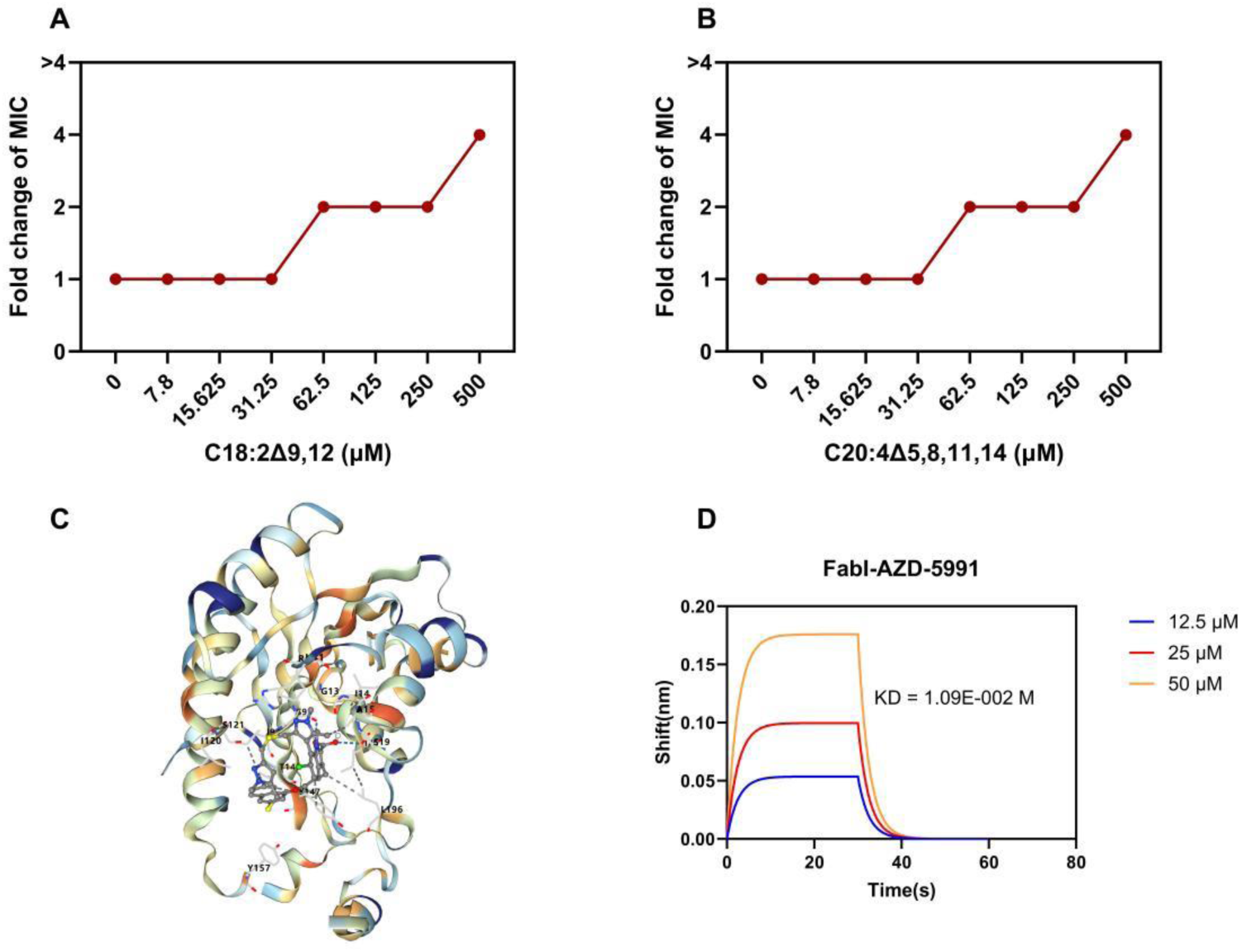
Effects of exogenous fatty acids on the antibacterial activity of AZD-5991. (A-B) Addition of exogenous fatty acids linoleic acid (C18:2Δ9,12) and arachidonic acid (C20:4Δ5,8,11,14) could significantly increase the MIC of AZD-5991 against S. *aureus* SA113. (C) Predicted molecular docking of AZD-5991 in the FabI binding pocket. (D) Kinetic analysis by BLI of the binding of AZD-5991 to FabI.

## Discussion

Numerous human tumors have elevated Mcl-1 expression, which is strongly linked to drug resistance, chemotherapy, and disease recurrence [13]. AZD-5991 is a kind of Mcl-1 inhibitor that activates the Bak-dependent mitochondrial apoptotic pathway by directly binding to Mcl-1 and causing cancer cells, including myeloma and acute myeloid leukemia, to undergo rapid apoptosis [9]. Here, our results indicated the excellent antibacterial activity of AZD-5991 against clinically isolated *S. aureus* and S*.epidermidis*. Compared to vancomycin, the time-killing dynamic curve assay showed AZD-5991 had more pronounced bactericidal action, particularly against MRSA. In addition, a major virulence factor that contributes to persistent infections is the production of biofilms by *S. aureus*, and only a small number of conventional antibiotics are effective against biofilms [15]. Significantly, AZD-5991 inhibited the production of *S. aureus* (both MSSA and MRSA) biofilms strongly. These findings suggest that AZD-5991 might be considered an antimicrobial agent in the clinical management of *S. aureus* infections.

Multidrug-resistant gram-positive pathogenic bacteria can commonly emerge as a primary cause of hospital-acquired or community-acquired infections [16]. Linezolid, a novel antibiotic licensed in 2000 for the antimicrobial treatment of infections brought on by MRSA and VAN-resistant enterococci (VRE), has been extensively documented in published research [17]. Even to this day, most gram-positive bacteria isolated from clinical samples around the world are still resistant to linezolid [18]. Nonetheless, our results reveal that AZD-5991 can inhibit the growth of linezolid-resistant *E. faecalis* and *E. faecium* isolates.

To date, only a single Phase I clinical trial (NCT03218683) has been conducted to evaluate the efficacy of AZD-5991 in patients with hematologic malignancies [19]. Consequently, studies on the pharmacokinetics, safety, and tolerability of this drug remain limited to preclinical animal experiments. A study using a mouse model indicated that a single intravenous injection of AZD-5991 at doses ranging from 10 to 100 mg/kg resulted in a plasma concentration of approximately 3 ng/mL at 24 hours post-administration. Furthermore, AZD-5991 demonstrated good tolerability at all tested dose levels, with no significant weight loss observed [9]. Therefore, at safe dose levels, the plasma concentrations of AZD-5991 can exceed the MIC values for most Gram-positive bacterial strains, suggesting its promising potential for clinical application. Additionally, the cytotoxicity of AZD-5991 against host cells was evaluated using CCK-8 assays in various cell lines, including LX-2, HepG2, HCT116, A549, and 293T. Importantly, these cytotoxic concentrations are significantly higher than the MIC values of AZD-5991 against *S*. *aureus* strains. This finding suggests a favorable therapeutic index for AZD-5991, highlighting its potential superior safety profile for future clinical applications. Totally to say, AZD-5991 with a high plasma concentration of over MIC might be toxic to the host, whereas oral administration of this chemical might have excellent effects for gastrointestinal infection by the effective concentration of AZD-5991 accumulation.

Furthermore, proteomic analysis of AZD-5991-treated *S*. *aureus* revealed significant changes in the expression levels of proteins involved in metabolic and transport pathways. Specifically, downregulated proteins, such as delta-hemolysin and ferric uptake regulation protein, have been implicated in bacterial virulence and iron homeostasis, which are crucial for *S*. *aureus* survival under stress conditions [20–21]. Conversely, upregulated proteins, including ketol-acid reductoisomerase (IlvC) and ribosomal subunit proteins, suggest an adaptive response aimed at counteracting cellular damage and sustaining metabolic activity [22–23]. The enrichment of pathways such as amino acid biosynthesis and pyrimidine metabolism underscores the bacterium’s attempts to restore cellular equilibrium following membrane perturbation induced by AZD-5991. The checkerboard assay findings further reinforce the role of phospholipids in modulating AZD-5991 activity. This phenomenon has been similarly reported with daptomycin, where phospholipids neutralized the antibiotic’s activity by binding to its hydrophobic tail [24]. However, unlike daptomycin, our BLI kinetic analysis showed no direct binding between AZD-5991 and cardiolipin, suggesting that the mechanism of AZD-5991 is more nuanced and may involve indirect disruption of membrane-associated processes rather than direct lipid binding. Interestingly, despite the substantial impact of AZD-5991 on membrane permeability and fatty acid metabolism, transmission electron microscopy (TEM) did not reveal significant morphological changes in treated cells. This observation supports the hypothesis that AZD-5991 induces localized damage to the cytoplasmic membrane without catastrophic disintegration, enabling solute leakage and bacterial death. Similar mechanisms have been described for other membrane-targeting agents, such as polymyxins, which create transient pores in the membrane rather than causing wholesale lysis [25].

Determining the precise mechanism of antibacterial compounds not only helps to improve the therapeutic benefits of outstanding derivatives but also offers new targets for the development and modification of antibacterial compound research [26]. The genetic mutations in the AZD-5991-induced tolerant *S*. *aureus* were detected by whole-genome sequencing and worthy of our attention, fatty acid efflux pump transcriptional regulator FarR, was proved to have a non-synonymous mutation. This finding implied that FarR gene mutations might play a significant role in the mechanism by which AZD-5991 causes resistance in *Staphylococcus aureus*. Therefore, mutations in FarR and its regulatory pathways likely suggest a mechanism by which AZD-5991 exerts selective pressure on *S*. *aureus*, leading to the emergence of resistant strains. As a key regulator of fatty acid metabolism, FarR plays a central role in controlling the synthesis and efflux of long-chain fatty acids, essential components of membrane phospholipids [30]. The observed mutation in FarR, coupled with the impact of exogenous fatty acids on the MIC values of AZD-5991, strongly supports the hypothesis that AZD-5991 disrupts bacterial membrane function by targeting fatty acid-related pathways. In our study, the significant reduction in antibacterial efficacy upon supplementation with unsaturated long-chain fatty acids, such as linoleic acid and arachidonic acid, indicates that these lipids may competitively inhibit AZD-5991’s interaction with its target sites. Mounting evidence has demonstrated the critical role of FabI in fatty acid-related pathways, which is closely correlated with FarR protein functions. FabI protein has been considered as the most important potential target of fatty acid-related pathways for the development of antimicrobial agents. Several antimicrobial agents, such as triclosan and cerulenin, have been reported to interact with FabI protein[31–32]. Therefore, we hypothesized that AZD5991 might target FabI for inhibiting bacterial growth. Interestingly, the results of the BLI kinetic studies revealed a binding interaction between AZD-5991 and FabI, a key enzyme in the fatty acid biosynthesis pathway. FabI inhibitors have been widely studied as promising antibacterial agents due to their ability to disrupt the synthesis of membrane phospholipids, causing bacteriostatic or bactericidal effects [33]. The compensatory mutations in FarR and the observed elevation in MIC values under exogenous fatty acid supplementation suggest that *S*. *aureus* may adapt to AZD-5991 exposure by altering fatty acid metabolism. This adaptation likely involves upregulation of alternative pathways or efflux mechanisms to maintain membrane integrity, a phenomenon previously documented in resistant strains of *S*. *aureus* exposed to lipid-targeting agents [34].

In conclusion, our findings suggest that AZD-5991 has antibacterial action against Gram-positive bacteria, efficiently destroying bacteria within mature biofilms and greatly reducing the production of biofilms. AZD-5991 exerts its antibacterial activity through a multifaceted mechanism involving the disruption of membrane permeability, interference with fatty acid-related pathways, and global metabolic reprogramming in *S*. *aureus*. The emergence of resistant mutants with compensatory mutations in FarR highlights the adaptability of *S*. *aureus* and underscores the importance of exploring combination therapies or targeting secondary pathways to mitigate resistance development. Further studies, such as structural analysis of AZD-5991-FabI interactions and the exploration of other potential targets, are warranted to fully elucidate the antibacterial mechanism of AZD-5991 and guide the development of next-generation derivatives.

## Materials and Methods

### Bacterial isolates and growth conditions

Strains of *S*. *aureus* (MRSA, n = 20 and MSSA, n = 20), *E*. *faecalis*(n = 21), *E*. *faecium*(n = 19), and *S*. *Epidermids* (n = 22) were isolated from different clinical specimens of individual patients at Shenzhen Nanshan People’s Hospital from 2011 to 2015. The isolates were sourced from urine, secretions, blood, bile, sputum, pus, etc. and were identified with standard methods using a VITEK 2 system (BioMérieux, Marcy l’ Etoile, France). *E*. *faecalis* ATCC29212 and OG1RF (ATCC 47077), and *S*. *aureus* USA300 were tested as quality control strains. Strains of *S*. *aureus* and *E*. *faecalis* were cultured in trypticsoy broth (TSB), with or without added 0.5% glucose (TSBG). Bacteria were routinely grown at 37°C and 220 rpm. The identified strains were stored at 80°C in 15% glycerol containing with TSB for further investigation.

### MIC assays

The MICs of antimicrobial agents were detected by the broth microdilution method in cation-adjusted Mueller–Hinton broth (CAMHB) according to the Clinical and Laboratory Standards Institute guidelines (CLSI M100, 29^th^ ed.). AZD-5991, ampicillin (AMP), vancomycin (VAN), and daptomycin (DAP) were purchased from MCE (Princeton, United States).

### Bacterial growth curve assay

Overnight cultures of *S*. *aureus* and *E*. *faecalis* were sub-cultured (1:200) in fresh TSB medium and AZD-5991 were coubling diluted (from 1 × MIC to 1/16 × MIC) by TSB medium. After diluted, 150 μL drug fluid and 150 μL bacterial fluid were added to growth curve device (Bioscreen C co., Piscataway, USA) in triplicate. The bacterial solution without AZD-5991 was used as positive control and OD_600_ value was read every half an hour and monitored continuously for 16 hours. The growth curve was drawn and the experiment was repeated three times.

### Time-kill dynamic curve assay

Overnight cultures of *S*. *aureus* YUSA145 and CHS101 were sub-cultured (1:200) in fresh TSB medium and incubated at 220 rpm at 37℃ until logarithmic growth period. The medium was adjusted by sterile normal saline until the bacterial suspension density was equal to a 0.5 McFarland standard (∼ 1.0 × 10^8^ CFU/mL) and then diluted to 1:100 into CAMHB medium. 1 × MIC, 2 × MIC, 4 × MIC, 8 × MIC AZD-5991 and 4 × MIC vancomycin were added and the medium was shaking-cultured at 220 rpm at 37℃. Aliquots (1 mL) were removed from the medium at different time points (2, 4, 6 and 24 h) and were centrifuged for 5 minutes and then discarded the supernatant. Bacterial solution was washed twice by 1mL 0.9% sterile NaCl and then resuspended. After multiple dilution, 100 μL of bacteria were evenly coated on TSA plate and cultured at 37℃ for 24 h. This experiment was repeated three times.

### Biofilm assay

We used microplate semi-quantitative method to observe the inhibitory effect of AZD-5991 on biofilm formation of *S*. *aureus* for 24 hours. The steps were as follows: overnight cultures of *S*. *aureus* strainswere sub-cultured (1:200) in fresh TSB medium and incubated until logarithmic growth period. After adjusting by sterile normal saline until the bacterial suspension density was equal to a 0.5 McFarland standard (∼1.0 × 10^8^ CFU/mL), the bacteria were diluted 1:100 by fresh TSBG (trypticsoybroth plus 0.25% glucose) and then inoculated onto 96-well microtiter plates (100μL/well). AZD-5991 (100μL/well) were added as well and mixed well till the concentration came to 1/2, 1/4 and 1/8 × MIC. Microplates were incubated at 37 ℃ for 24 h, and wells with no derivatives were used as positive control. Then bacterial solution was removed and wells were washed by deionized water for three times. After drying at room temperature, adherent biofilms were fixed with 95% methanol and stained with 1% crystal violet (100μL) for 20 min and washed. The optical density at 570 nm (OD_570_) was then tested by micro plate spectrophotometer. And this experiment was repeated three times.

### Cell proliferation assay and cytotoxicity assay

A cell count kit-8 (CCK-8 Beyotime, China) was used in the cytotoxicity test.100 μL of different cell suspension (LX-2, HepG2, HCT116, A549 and 293T cells; 5000 cells/well) were dispensed in a 96-well plate. After that, various concentrations of AZD-5991 from 0.78 μM to 200 μM (10 μL) were added to the plate. The plate was then incubated for an appropriate length of time (e.g. 6,12,24 or 48 hours) in the incubator. After incubation, 10 μL of CCK-8 solution was added to each well of the plate. Next, the plate was incubated for 1-4 hours in the incubator and measured the absorbance at 450 nm by a microplate reader.

### Membrane permeability and depolarization assay

*S*. *aureus* SA113 were cultured overnight in TSB medium at 37℃. Subsequently, 10^7^ CFU/mL bacteria were added to HEPES medium (5 mM HEPES, 20 mM glucose, pH 7.4) and incubated with 1 mM Propidium Iodide (PI) (excitation l = 217 nm, emission l = 340 nm) or 1 mM diSC3(5) (excitation l = 622 nm, emission l = 673 nm) for 2 h, then KCl was added to a final concentration of 0.1 M to balance cytoplasmic and external K^+^ concentrations. The bacteria were placed in a 96-well plate for a moment until the fluorescence intensity remained stable, then AZD-5991 was added to a final concentration of 1 ×, 2× and 4 × MIC and the fluorescence intensity was recorded continuously by a high-content instrument. Samples without AZD-5991 and with 0.1% Triton X-100 were used as the negative and positive control, respectively.

### TEM imaging assay

Overnight cultures of *S*. *aureus* YUSA145 were diluted 1:100 in TSB medium to a cell density of 10^7^ CFU/mL and cultured at 37 ℃ for 4 h. AZD-5991 (8 × MIC) was added to cells and the mixture was incubated at 37℃ for 1 h. DMSO treatment was considered as a negative control. Then, the cells were collected by centrifugation at 4,000 rpm for 5 min, and then washed with PBS and fixed overnight with fixative (4% paraformaldehyde and 0.1 M phosphate buffer) at 4 ℃. After discarding the supernatant of fixed cells, 0.1 M PB (pH=7.4) was added, and the cells were re-suspended and washed three times in PB for 3 min. The sample was then dehydrated with gradient ethanol (30, 50, 70, 80, 95, 100%) for 20 min. Next, the sample was treated with acetone and EMBed 812 (1:1) for 2hat at 37℃, followed by acetone and EMBed 812 (1:2) overnight at 37℃, and then EMBed 812 for 6h at 37℃ The embedding models with resin and samples were polymerized by heating to 65℃ for more than 48 h. The resin blocks were cut to 60–80 nm thickness on a LEICA EM UC7 ultrathin slicer, and the sample was fished out onto 50 mesh copper grids with formvar film. After being staining with 2.6% lead citrate for 8 min, and then rinsed three times with ultra-pure water. The copper grids were dried and placed into the grids board, and then the samples were observed under a transmission electron microscope (TEM, HITACHI HT 7800).

### In vitro selection of S. *aureus* exhibiting AZD-5991-induced resistance

*S*. *aureus* isolate YUSA145 with MICs of 3.125 μM were subjected to induce AZD-5991-resistance in vitro. The isolates were subcultured serially in medium supplied with increasing AZD-5991 concentrations from 1/2 ×, 1 ×, 2 ×, 4 ×, and 8 × MIC, and successively passaged four passages at each concentration. Isolates at each concentration were collected and cultured three continuous generations without AZD-5991 for further studies. And isolates exhibiting AZD-5991-induced resistance with the 8 × MIC were employed to explore genetic variability by whole-genome sequencing.

### Bacterial whole-genome sequencing

Chromosomal DNA of AZD-5991-induced resistant *S*. *aureus* strain YUSA145 was prepared for whole-genome sequencing. Nextera libraries construction and whole genome sequencing was performed on Illumina HiSeq sequencing platform by Novogene Co. Ltd., (Beijing, China). The sequencing reads were mapped against the *S*. *aureus* strain SA113 reference genome in bwa-mem software (v0.7.5a)2 with standard parameters. Single nucleotide polymorphisms and indels in resistant strains SA113 were displayed using MUMmer (version 3.23). The data were deposited in the National Center for Biotechnology Information (NCBI) with accession number PRJNA1110364 (NCBI: https://www.ncbi.nlm.nih.gov/bioproject/1110364).

### Sample preparation for quantitative proteomics

*S. aureus* SA113 at exponential growth phase (OD_600_ of 0.5) was added with AZD-5991 to final concentrations that corresponded to 1/2 × MIC. The sham group was treated with DMSO. Each group was performed with three biological replicates. The cultures were then incubated at 37°C for 2 h on a shaker at 200 rpm. After that, the bacteria were harvested by centrifugation at 5000 g for 10 min at 4 ℃. After washed with cold PBS three times, the cell pellets were suspended in RIPA lysis buffer (1% Triton X-100, 1% deoxycholate, 0.1% SDS) with complete protease inhibitor cocktail (catalog No. 05892970001, Roche, Basel, Switzerland). The suspension was then subjected to three rounds of homogenization with glass beads (diameter 0.1 mm) and centrifuged at 12000 g for 20 min at 4 °C, and the supernatants were collected for protein concentration determination and subsequent quantitative proteomics. Pierce Micro BCA Protein Assay Kit (catalog No. 23227, Thermo Fisher Scientific, MA, USA) was used to determine the protein concentration. 100 μg of extracted protein was reduced with 10 mM DTT (Sigma-Aldrich Co., St. Louis, MO) for 1 h at 70 °C, followed by alkylation using 50 mM iodoacetamide (IAA, Sigma-Aldrich) for 15 min at room temperature in the dark. The samples were then desalted and buffer-changed three times with 100 μL 0.5 M ammonium bicarbonate by using Amicon Ultra Centrifugal Filters (10 kDa cutoff; Millipore, Billerica, MA). The proteins were digested with trypsin (Promega, Madison, WI) at a ratio of 1:50 at 37 ℃ overnight. They were then lyophilized and stored at −80 ℃.

### Nano LC-MS/MS Analysis for quantitative proteomics

Samples were reconstituted in 30 μL of 0.1% formic acid, among which 4 μL of each sample was injected onto an LC system consisting of an UltiMate 3000 RSLC nano system and a C18 precolumn (100 μm × 20 mm, Acclaim PepMap 100 C18, 3 μm), followed by separation using a C18 tip column (75 μm × 250 mm, Acclaim PepMap RSLC, 2 μm). The mobile phases A and B were composed of 0.1% formic acid and 80% acetonitrile in 0.1% formic acid, respectively. The elution system started with 5% B for the first 5 min, followed by a linear gradient from 5% B to 38% B in the next 85 min and from 38% B to 95% B in the next 2 min, maintained at 95% B for another 3 min at a flow rate of 300 nL/min. The column was coupled to Q Exactive Plus mass spectrometer equipped with the Nano spray ionization (NSI) interface. MS1 scans were acquired over a mass range of 300-1500 m/z with a resolution of 70,000 and the corresponding MS2 spectra were acquired at a resolution of 17500, collected for maximally 50 ms. All multiply charged ions were used to trigger MS-MS scans followed by a dynamic exclusion for 30 s. Singly charged precursor ions and ions of undefinable charged states were excluded from fragmentation.

### Bioinformatics Analysis for quantitative proteomics

The protein identification and quantification were performed using Proteome Discoverer 2.4 base with the Sequest HT against the Uniprot proteome of Enterococcus faecalis EnGen0311 (Strain: TX0635). A 2-fold cutoff value was applied to determine up regulated and down regulated proteins in addition to a p-value of less than 0.05 in at least two technical replicates. The differentially expressed proteins were uploaded into the OMICSBEAN database (http://www.omicsbean.com) for GO (gene ontology) annotation, including biological process, cellular component, molecular function, and KEGG pathway analysis. The PPI networks were analyzed by using this web-based tool OMICSBEAN.

### Antibacterial activity of AZD-5991 with phospholipids and fatty acids

Methanol was used to dissolve a variety of phospholipids and fatty acids, including arachidonic acid (A5837, Sigma-Aldrich, USA). The chequerboard method, as previously reported [35], was used to assess the impact of these phospholipids (concentrations ranging from 8 to 128 μg/mL) and fatty acids (concentrations ranging from 7.8 to 500 μM) on the MIC values of AZD-5991 in TSB medium.

### Predicted molecular docking

FabI (Uniprot ID: Q2FVQ3)’s molecular structure PDB file was generated using AlphaFoldDB [36]. The Protein Preparation Wizard feature of the Schrödinger software was used to perform hydrogenation, replace missing residues, and optimise structure and water molecules. The molecular 3D structure file for AZD-5991 (CAS No. HY-101533) was retrieved from PubChem. Molecular docking was used to establish the binding architecture of the FabI protein and AZD-5991. In conclusion, structure-based cavity detection was utilised to choose the optimal binding pocket of the FabI protein, and AutoDock Vina was used to investigate the optimum binding location of AZD-5991 based on the Vina score (kcal/mol) in the pocket [37].

### Biolayer interferometry assay (BLI)

Using a biolayer interferometry (BLI) test with Gatorprime equipment (Gator Bio, San Francisco, USA), the binding affinities between AZD-5991 and FabI were evaluated using biotinylated cardiolipin (L-C16B, Echelon Biosciences, USA). The tips of streptavidin biosensors were immobilised using biotin-labeled FabI after being conditioned with kinetic buffer (PBS, 0.05% bovine serum albumin, and 0.01% Tween 20). The biosensors were then treated with AZD-5991 at different concentrations. In order to account for nonspecific binding and signal variability, duplicate sets of biosensors were used as background controls and incubated in protein-free buffer. Following established procedures, the tests were carried out in 96-well black plates at 30 ℃ with a total volume of 300 μL per well. Gatorprime software was used for data analysis, and a double reference subtraction method was used to accurately determine binding kinetics.

### Quantification and statistical analysis

Statistical analysis was performed using GraphPad Prism software (version 9.0). The Log-rank (Mantel-Cox) test was utilized to assess survival rates. Data are presented as mean ± SD, with a p-value of <0.05 deemed statistically significant (*p < 0.05, **p < 0.01, ***p < 0.001). Most experiments were conducted at least thrice, yielding similar results.

## Author Contributions

S.H. and Z.C. conceived and designed the project. Y.T., Z.X., H.D., and X.Y. conducted the experiments. The other authors analyzed experimental results. Y.T. wrote the manuscript. Z.Y. and Z.C. revised the manuscript. S.H., Z.C., and T.H. acquired the funding. All authors have read and approved the manuscript.

## Notes

The authors declare that they have no conflicts of interest.

## ACKNOWLEDGMENT

The authors thank Weiguang Pan (Department of Laboratory Medicine, Shenzhen Nanshan People’s Hospital, Shenzhen 518052, China) for helping identify and preserve the bacterial isolates.

## ABBREVIATIONS

*E. faecalis*: *Enterococcus faecalis*
*S. aureus*: *Staphylococcus aureus*
VRE: Vancomycin-resistant enterococci
MDR: Multidrug-resistant
MRSA: Methicillin-resistant *Staphylococcus aureus*
MSSA: Methicillin-sensitive *Staphylococcus aureus*

## REFERENCES

[1] Bhowmik A. Role of Diagnostic procedures in managing human Bacterial infections: A comprehensive overview. Arch Hematol Case Rep Rev. 2023, 8 (1), 008–019.

[2] Douglas E J A, Wulandari S W, Lovell S D, Laabei M. Novel antimicrobial strategies to treat multi-drug resistant Staphylococcus aureus infections. Microb Biotechnol. 2023, 16 (7), 1456–1474.

[3] Hu F P, Guo Y, Wang F, Zhu D M. Chinet 2021 surveillance of bacterial resistance in china. Chinese Journal of Infection and Chemotherapy. 2022, 22 (5), 521–530.

[4] Abebe A A, Birhanu A G. Methicillin resistant Staphylococcus aureus: molecular mechanisms underlying drug resistance development and novel strategies to Combat. Infect Drug Resist. 2023, 16, 7641–7662.

[5] Kamble E, Pardesi K. Antibiotic tolerance in biofilm and stationary-phase planktonic cells of Staphylococcus aureus. Microb. Drug Resist. 2021, 27 (1), 3–12.

[6] Parastan R, Kargar M, Solhjoo K, Kafilzadeh F. Staphylococcus aureus biofilms: Structures, antibiotic resistance, inhibition, and vaccines. Gene Rep. 2020, 20, 100739.

[7] Ulloa E R, Singh K V, Geriak M, Haddad F, Murray B E, Nizet V, Sakoulas G. Cefazolin and ertapenem salvage therapy rapidly clears persistent methicillin-susceptible Staphylococcus aureus bacteremia. Clin Infect Dis. 2020, 71 (6), 1413–1418.

[8] Xu L, She P, Chen L, Li S, Zhou L, Hussain Z, Liu Y, Wu Y. Repurposing candesartan cilexetil as antibacterial agent for MRSA infection. Front Microbiol. 2021, 12, 688772.

[9] Tron A E, Belmonte M A, Adam A, Aquila B M, Boise L H, Chiarparin E, Cidado J, Embrey K J, Gangl E, Gibbons F D, Gregory G P, Hargreaves D, Hendricks J A, Johannes J W, Johnstone R W, Kazmirski S L, Kettle J G, Lamb M L, Matulis S M, Nooka A K, Packer M J, Peng B, Rawlins P B, Robbins D W, Schuller A G, Su N, Yang W, Ye Q, Zheng X, Secrist J P, Clark E A, Wilson D M, Fawell S E, Hird A W. Discovery of Mcl-1-specific inhibitor AZD5991 and preclinical activity in multiple myeloma and acute myeloid leukemia. Nat Commun. 2018, 9 (1), 5341.

[10] Jennings J A, Beenken K E, Skinner R A, Meeker D G, Smeltzer M S, Haggard W O, Troxel K S. Antibiotic-loaded phosphatidylcholine inhibits staphylococcal bone infection. World J Orthop. 2016, 7 (8), 467.

[11] Kopiasz R J, Rukasz A, Chreptowicz K, Podgórski R, Kuźmińska A, Mierzejewska J, Tomaszewski W, Ciach T, Jańczewski D. Influence of lipid bilayer composition on the activity of antimicrobial quaternary ammonium ionenes, the interplay of intrinsic lipid curvature and polymer hydrophobicity, the role of cardiolipin. Colloid Surface B. SCIE. 2021, 207, 112016.

[12] Nicola N. Can you teach old drugs new tricks?. Nature. 2016, 534 (7607), 314–316.

[13] Wuillème-Toumi S, Robillard N, Gomez P, Moreau P, Le Gouill S, Avet-Loiseau H, Harousseau J L, Amiot M, Bataille R. Mcl-1 is overexpressed in multiple myeloma and associated with relapse and shorter survival. Leukemia. 2005, 19 (7), 1248–1252.

[14] Sakoulas G, Moise-Broder P A, Schentag J, Forrest A, Moellering R C, Eliopoulos G M. Relationship of mic and bactericidal activity to efficacy of vancomycin for treatment of methicillin-resistant staphylococcus aureus bacteremia. J Clin Microbiol. 2004, 42 (6), 2398–402.

[15] Shree P, Singh C K, Sodhi K K, Surya J N, Singh D K. Biofilms: Understanding the structure and contribution towards bacterial resistance in antibiotics. Medicine Microecol. 2023, 16, 100084.

[16] Cornejo-Juárez P, Vilar-Compte D, Pérez-Jiménez C, Ñamendys-Silva S A, Sandoval-Hernández S, Volkow-Fernández P. The impact of hospital-acquired infections with multidrug-resistant bacteria in an oncology intensive care unit. Int J Infect Dis. 2015, 31, 31–34.

[17] Gonzales R D, Schreckenberger P C, Graham M B, Kelkar S, DenBesten K, Quinn J P. Infections due to vancomycin-resistant Enterococcus faecium resistant to linezolid. Lancet. 2001, 357 (9263), 1179.

[18] Jubeh B, Breijyeh Z, Karaman R. Resistance of gram-positive bacteria to current antibacterial agents and overcoming approaches. Molecules. 2020, 25 (12), 2888.

[19] Desai P, Lonial S, Cashen A, Kamdar M, Flinn I, O’Brien S, Garcia J S, Korde N, Moslehi J, Wey M, Cheung P, Sharma S, Olabode D, Chen H, Syed F A, Liu M, Saeh J, Andrade-Campos M, Kadia T M, Blachly J S. A phase 1 first-in-human study of the MCL-1 inhibitor AZD5991 in patients with relapsed/refractory hematologic malignancies. Clin Cancer Res. 2024, 30 (21): 4844–4855.

[20] Chen J, Lv Y, Shang W, Yang Y, Wang Y, Hu Z, Huang X, Zhang R, Yuan J, Huang J, Rao X. Loaded delta-hemolysin shapes the properties of Staphylococcus aureus membrane vesicles. Front Microbiol. 2023, 14, 1254367.

[21] Fontenot C R, Tasnim H, Valdes K A, Popescu C V, Ding H. Ferric uptake regulator (Fur) reversibly binds a [2Fe-2S] cluster to sense intracellular iron homeostasis in Escherichia coli. J Biol Chem. 2020, 295 (46), 15454–15463.

[22] Wong S H, Lonhienne T G, Winzor D J, Schenk G, Guddat L W. Bacterial and plant ketol-acid reductoisomerases have different mechanisms of induced fit during the catalytic cycle. J Mol Biol. 2012, 424 (3-4), 168–179.

[23] Akanuma G, Nanamiya H, Natori Y, Yano K, Suzuki S, Omata S, Ishizuka M, Sekine Y, Kawamura F. Inactivation of ribosomal protein genes in Bacillus subtilis reveals importance of each ribosomal protein for cell proliferation and cell differentiation. J Bacteriol. 2012, 194 (22), 6282–6291.

[24] Ramamoorthy A. NMR structural studies on membrane proteins. BBA Bioenergetics. 2007, 1768 (12), 2947–2948.

[25] Mohapatra S S, Dwibedy S K, Padhy I. Polymyxins, the last-resort antibiotics: Mode of action, resistance emergence, and potential solutions. J Biosciences. 2021, 46 (3), 85.

[26] Vaou N, Stavropoulou E, Voidarou C, Tsigalou C, Bezirtzoglou E. Towards advances in medicinal plant antimicrobial activity: A review study on challenges and future perspectives. Microorganisms. 2021, 9 (10), 2041.

[27] Nguyen M T, Saising J, Tribelli P M, Nega M, Diene S M, François P, Schrenzel J, Spröer C, Bunk B, Ebner P, Hertlein T, Kumari N, Härtner T, Wistuba D, Voravuthikunchai S P, Mäder U, Ohlsen K, Götz F. Inactivation of farR causes high rhodomyrtone resistance and increased pathogenicity in Staphylococcus aureus. Front Microbiol. 2019, 10, 1157.

[28] Limsuwan S, Trip E N, Kouwen T R, Piersma S, Hiranrat A, Mahabusarakam W, Voravuthikunchai S P, Dijl J M, Kayser O. Rhodomyrtone: a new candidate as natural antibacterial drug from Rhodomyrtus tomentosa. Phytomedicine. 2009, 16 (6-7), 645–651.

[29] Alnaseri H, Kuiack R C, Ferguson K A, Schneider J E, Heinrichs D E, McGavin M J. DNA binding and sensor specificity of FarR, a novel TetR family regulator required for induction of the fatty acid efflux pump FarE in Staphylococcus aureus. J Bacteriol. 2019, 201 (3), 10–1128.

[30] Heath R J, Rubin J R, Holland D R, Zhang E, Snow M E, Rock C O. Mechanism of triclosan inhibition of bacterial fatty acid synthesis. J Biol Chem. 1999, 274 (16), 11110–11114.

[31] Singh V, Harinarayanan R. (p) ppGpp Buffers Cell Division When Membrane Fluidity Decreases in Escherichia coli. Mol Microbiol. 2024, 122 (6), 847–865.

[32] Payne D J, Miller W H, Berry V, Brosky J, Burgess W J, Chen E, DeWolf W E, Fosberry A P, Greenwood R, Head M S, Heerding D A, Janson C A, Jaworski D D, Keller P M, Manley P J, Moore T D, Newlander K A, Pearson S, Polizzi B J, Qiu X, Rittenhouse S F, Slater-Radosti C, Salyers K L, Seefeld M A, Smyth M G, Takata D T, Uzinskas I N, Vaidya K, Wallis N G, Winram S B, Yuan C C K, Huffman W F. Discovery of a novel and potent class of FabI-directed antibacterial agents. Antimicrob Agents CH. 2002, 46 (10), 3118–3124.

[33] Huang L, Matsuo M, Calderón C, Fan S H, Ammanath A V, Fu X, Li N, Luqman A, Ullrich M, Herrmann F, Maier M, Cheng A, Zhang F, Oesterhelt F, Lämmerhofer M, Götz F. Molecular basis of rhodomyrtone resistance in Staphylococcus aureus. Mbio. 2022, 13 (1), e03833–21.

[34] Chen J, Lv Y, Shang W, Yang Y, Wang Y, Hu Z, Huang X, Zhang R, Yuan J, Huang J, Rao X. Loaded delta-hemolysin shapes the properties of Staphylococcus aureus membrane vesicles. Front Microbiol. 2023, 14, 1254367.

[35] Lee A S, De Lencastre H, Garau J, Kluytmans J, Malhotra-Kumar S, Peschel A, Harbarth S. Methicillin-resistant Staphylococcus aureus. Nat Rev Dis Primers. 2018, 4 (1), 1–23.

[36] Jumper J, Evans R, Pritzel A, Green T, Figurnov M, Ronneberger O, Tunyasuvunakool K, Bates R, Žídek A, Potapenko A, Bridgland A, Meyer C, Kohl S A A, Ballard A J, Cowie A, Romera-Paredes B, Nikolov S, Jain R, Adler J, Back T, Petersen S, Reiman D, Clancy E, Zielinski M, Steinegger M, Pacholska M, Berghammer T, Bodenstein S, Sliver D, Vinyals O, Senior A W, Kavukcuoglu K, Kohli P, Hassabis D. Highly accurate protein structure prediction with AlphaFold. Nature. 2021, 596 (7873), 583–589.

[37] Liu Y, Yang X, Gan J, Chen S, Xiao Z X, Cao Y. CB-Dock2: Improved protein–ligand blind docking by integrating cavity detection, docking and homologous template fitting. Nucleic Acids Res. 2022, 50 (W1), W159–W164.

